# An annotation-overlap-flagged rare-disease gene-prioritisation benchmark and PMC index recipe

**DOI:** 10.64898/2026.08.30.748028

**Authors:** Johanna Angulo, Hector Espinos-Morato, Víctor Yeste

**Author notes:** Corresponding author: Hector Espinos-Morato.

## Abstract

Benchmarks for rare-disease gene prioritisation are assembled from published clinical cases. Those cases often come from the same publications used to build knowledge-base (“curated”) tools, so a curated tool can be scored on its own source literature. This resource makes that circularity measurable. We release a stratified benchmark of 1,047 rare-disease cases from the GA4GH Phenopacket Store v0.1.26. Each case pairs a Human Phenotype Ontology profile with a 50-gene candidate list (one causal gene, 49 distractors) and the causal-gene label, sampled across four operational MONDO-derived disease strata and issued in two case-paired variants: random distractors, and phenotype-similar distractors selected by HPO Resnik similarity. Two case-level metadata layers support fairer evaluation: a per-case flag recording whether a case’s source publication is cited in the HPO disease-annotation file, defining an overlap-absent subset (*n* = 282), and publication-recency strata. We also specify a deterministic, version-pinned recipe for a hybrid dense-plus-sparse retrieval index over *∼* 2.25 million PMC Open Access articles (52,777,395 chunks). The resource reports no tool comparisons.

## 1 Background & Summary

Rare diseases are individually uncommon but collectively affect an estimated 300 million people worldwide [1,2]. Despite international efforts to make a molecular diagnosis available for every rare genetic condition [3], roughly half of exome– or genome-sequenced cases remain undiagnosed after standard analysis [4]. This is often because current knowledge cannot confidently link the candidate variants that sequencing uncovers to the patient’s disease [5].

A central computational step is phenotype-driven gene prioritisation: given a structured description of a patient’s clinical features, rank the genes most likely to explain them. Features are recorded using the Human Phenotype Ontology (HPO), a standardised, hierarchically organised vocabulary of phenotypic abnormalities [6], so a patient is represented as a set of HPO terms. Tools such as Exomiser [7], LIRICAL [8] and Phen2Gene [9] score candidate genes against curated knowledge bases of disease–gene–phenotype associations. By construction, these tools cannot surface associations that have not yet been curated [10]. A newer class reasons over the primary literature using retrieval-augmented generation (RAG), which retrieves text passages and conditions a language model on them [11,12]. Biomedical examples include GeneGPT [13] and RAG systems for medical question answering [14], increasingly organised as multi-step workflows [15].

Comparing the two classes involves a specific circularity. Public rare-disease benchmarks are typically assembled from published, molecularly solved cases, a widely used source being the GA4GH Phenopacket Store [16,17]. Curated tools are built from that same literature: LIRICAL, for instance, computes likelihood ratios from the HPO disease-annotation file phenotype.hpoa, which records the publications each annotation was curated from. Where a benchmark case and a curated tool’s knowledge base trace to the same publication, the tool has been exposed to that case’s source. Such data leakage is a documented driver of over-optimistic results across machine-learning-based science [18]. Existing phenotype-prioritisation frameworks standardise how tools are scored [19], but they do not record which cases carry this exposure.

The resource has four components. The first is a stratified, version-pinned cohort of 1,047 cases with 50-gene candidate lists, sampled across four operational MONDO-derived strata (developmental, immunological, metabolic, neurological). Each case is supplied in two case-paired difficulty variants that regenerate from the same per-case seed: a standard variant, whose distractors are drawn at random from the HGNC-approved protein-coding set, and a hard variant, whose distractors are phenotype-similar genes selected by HPO Resnik semantic similarity. The second is a per-case annotation-overlap flag recording whether a case’s source publication is cited in phenotype.hpoa for the case’s own disease; we call the cases with no detected citation the overlap-absent subset (*n* = 282). The classification is reproducible from public inputs, but it does not guarantee that these cases are free from leakage (Section 5). The third is a set of publication-recency strata, supporting the separate question of generalisation to associations that post-date curation cycles. The fourth is a hybrid dense-plus-sparse retrieval index over the genetics-relevant PMC Open Access corpus (*∼*2.25 M articles; 52,777,395 chunks). Mixed per-article licences preclude redistributing the chunk text, so we release the index as a deterministic, version-pinned build recipe with content-addressed chunk identifiers and a chunk-set fingerprint.

Table 1 lists related resources for context.

**Table 1:** Related rare-disease prioritisation resources. The *Type* column distinguishes benchmark cohorts from evaluation frameworks and community challenge rounds: PhEval is a standardisation framework and the CAGI6 Rare Genomes Project round is a community challenge, so neither is a competing cohort and neither should be read as a cohort with missing features. The remaining columns record which features each resource provides; the overlap flag of the present resource is recomputable from public inputs against any phenotype.hpoa release.

| Resource | Type | Cases | Overlap flag | Difficulty variants | Retrieval sub-strate | Reproducibility guarantee | Licence |
| --- | --- | --- | --- | --- | --- | --- | --- |
| This resource | Cohort + index | 1,047, with 50-gene candidate lists | Yes | Yes (standard, hard; case-paired) | Yes (build recipe) | SHA-256-verified regeneration of both cohort files; content-addressed chunk identifiers | CC BY 4.0; AGPL-3.0 (code) |
| PhEval [19] | Evaluation framework | n/a (ships runners and example corpora) | No | No | No | Versioned runners and pinned corpora | Apache-2.0 |
| GA4GH Phenopacket Store [17] | Case corpus | 9,588 phenopackets; no candidate lists | No | No | No | Versioned, citable releases | CC BY 4.0 |
| CAGI6 Rare Genomes Project [20] | Community challenge | 65 families (30 test), with variant calls | No | No | No | Blinded challenge round | Controlled access |

The annotation-overlap procedure depends only on public inputs and can be recomputed against any phenotype.hpoa release, so it applies to other benchmarks assembled from published cases. In future work we intend to use the resource to evaluate an agentic-workflow, literature-based RAG prioritiser; evaluations of that kind also need to account for known failure modes of generative systems, notably hallucination [21]. Keeping the benchmark separable from the systems that use it allows others to adopt it independently.

## 2 Methods

This section describes how we assembled the benchmark cohort from public inputs, how we computed the two case-level metadata layers, and how we built the retrieval index. Every step is deterministic and version-pinned, so the whole resource can be regenerated from public data.

### 2.1 Benchmark cohort construction

We drew cases from GA4GH Phenopacket Store v0.1.26 (released 13 January 2026; [17]), which aggregates literature-curated rare-disease phenopackets with gene-level solved diagnoses. Of the 9,588 phenopackets loaded, inclusion required that three criteria be met.

The first criterion was exactly one distinct causal gene symbol across the phenopacket’s genomic interpretations, counting only interpretations whose interpretationStatus is CAUSATIVE, CONTRIBUTORY, or unspecified; cases with zero, or with two or more, distinct such genes were excluded. All 9,588 phenopackets in Store v0.1.26 carry a SOLVED progressStatus, so no additional filter on that field was applied.

The second criterion was *≥* 3 HPO terms (HPO v2026-02-16; [6]). The third was a mapping in the Mondo Disease Ontology (MONDO v2026-03-03; [22]) into one of four disease categories, each defined as the descendant closure (subclass closure, including the root itself) of fixed MONDO roots: *neurological* (MONDO:0005071, nervous system disorder), *metabolic* (MONDO:0005066, metabolic disease), *immunological* (MONDO:0005046, immune system disorder), and *developmental* (MONDO:0021147, inborn genetic disease, together with MONDO:0019118, developmental and epileptic encephalopathy).

HPO profiles comprise only observed (non-excluded) phenotypic features, deduplicated with document order preserved. Terms explicitly recorded as excluded in the phenopacket—that is, clinical features asserted to be absent—are not included in the profile.

#### Disease-scope exclusions

Two disease classes were excluded at the inclusion stage, independently of the four category roots: cases whose resolved MONDO identifiers fall under MONDO:0019042 (chromosomal disorder) or MONDO:0044970 (mitochondrial disease), each taken as its descendant closure.

Both classes sit awkwardly with a single-causal-gene, nuclear-protein-coding prioritisation task. Chromosomal disorders are typically caused by structural or copy-number events that span many genes rather than by a pathogenic variant in one gene [23], so the single-causal-gene labelling this benchmark assumes does not apply to them. Mitochondrial disease was excluded because its mitochondrial-genome-encoded causes lie outside the HGNC nuclear protein-coding candidate space from which distractors are drawn; applied at the disease-class level, the rule also removes nuclear-encoded mitochondrial disease [24]. Cases carrying no disease annotation were also excluded, since the annotation-overlap layer (Section 2.2) requires a disease identifier to join against. These exclusions restrict the cohort’s coverage and are revisited under Limitations (Section 5).

The first two criteria, together with the disease-scope exclusions, retained 6,382 cases. Of the 3,206 cases dropped, 1,155 had fewer than three HPO terms, 69 had no single ascertained causal gene, none had two or more, none lacked a disease annotation, and 1,982 fell under the chromosomal-disorder or mitochondrial-disease roots.

The third criterion then produced an eligible pool of 4,670 cases: 464 developmental, 390 immunological, 672 metabolic, and 3,144 neurological.

#### Category assignment and how to read it

The four descendant closures are not disjoint, so we resolve overlaps with an explicit rule: we assign each case to the *first* matching category in the fixed priority order neurological *>* metabolic *>* immunological *>* developmental, and retain the full list of matched categories per case for audit. Phenopacket diseases are cited by OMIM identifier, so we resolve each to MONDO through MONDO’s own cross-reference index, inverted once per run. Where more than one MONDO term declares the same cross-reference, we take the first in ontology traversal order. This is deterministic for a pinned MONDO release, but not necessarily the semantically preferred term.

The four categories are not parallel clinical domains. Three are body-system roots, whereas MONDO:0021147 (inborn genetic disease) subsumes most rare monogenic conditions, so the *developmental* stratum is a residual one: inborn genetic disease not already assigned to a body-system category. The strata spread the cohort across broad disease areas so that no single area dominates evaluation; they are not a clinical taxonomy, and per-category results should be read accordingly. We excluded cases matching none of the four roots (Figure 1).

**Figure 1:**
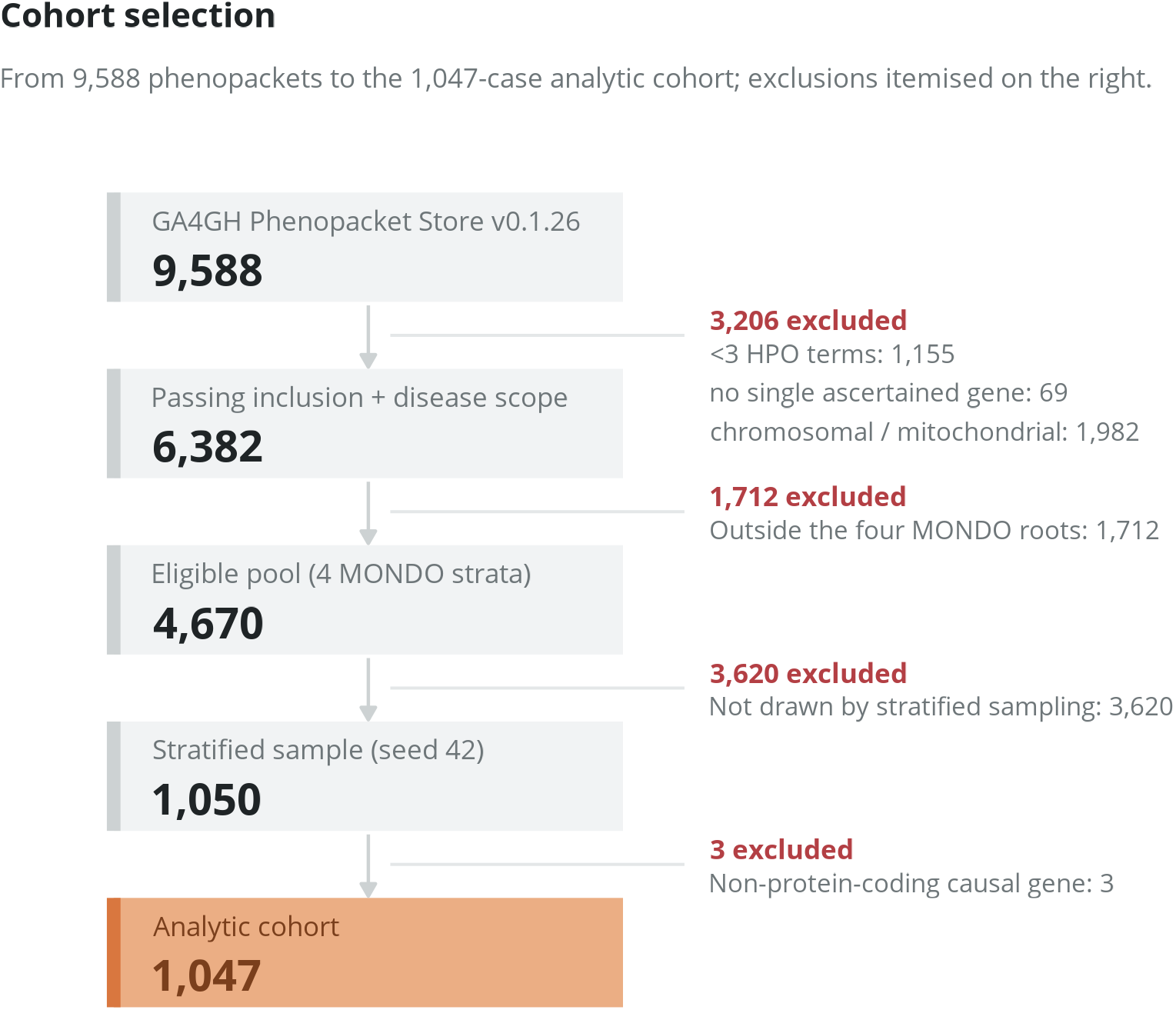
CONSORT-style cohort selection flow. GA4GH Phenopacket Store v0.1.26 is screened down to the final analytic cohort (*n* = 1,047) across four MONDO supercategories (developmental, immunological, metabolic, neurological). Exclusions are itemised, as is the downstream annotation-overlap stratification into the overlap-present and overlap-absent subsets.

The eligible pool is dominated by neurological cases, so a simple proportional sample would leave the smallest category (immunological) too sparsely represented for stable per-category evaluation. We therefore drew a disproportionate (oversampled) stratified sample of 1,050 cases with seed 42: 250 each from the developmental, metabolic, and neurological pools, and 300 from the immunological pool. Oversampling a rate-limiting stratum in this way is a standard sampling design: it protects the precision of estimates for the small subgroup, while stratum-weighted estimates remain unbiased for the protein-coding subset of the eligible pool once the design weights are applied [25]. Because we sampled the four strata at very different rates (Table 2), unweighted pooling over the cohort estimates a quantity defined by the sampling design rather than by the eligible population. We release the inclusion probabilities so that either quantity can be computed explicitly.

**Table 2:** Sampling design by disease category. The inclusion probability is the number *drawn* divided by the eligible pool, since the draw defines the design, and the design weight is its reciprocal. We removed three neurological cases after sampling because their causal genes are not protein-coding; the removal did not condition on the draw, so the design weight for a retained neurological case is unchanged at 3,144*/*250 *≈* 12.58. The analytic cohort therefore represents the protein-coding subset of each eligible pool, whose neurological size we estimate at 247 *×* (3,144*/*250) *≈* 3,106.

| Category | Eligible pool | Drawn | Analytic cohort | Inclusion prob. | Design weight |
| --- | --- | --- | --- | --- | --- |
| Developmental | 464 | 250 | 250 | 0.539 | 1.86 |
| Immunological | 390 | 300 | 300 | 0.769 | 1.30 |
| Metabolic | 672 | 250 | 250 | 0.372 | 2.69 |
| Neurological | 3,144 | 250 | 247 | 0.0795 | 12.58 |
| **Total** | **4,670** | **1,050** | **1,047** | — | — |

For each sampled case we counted the distinct PMC OA articles among the top-100 hybrid-retrieval hits for the causal-gene symbol; any case with fewer than five was to be replaced by a fresh case from the same category. All 1,050 sampled cases cleared the threshold (initial_fail = 0, replacements_made = 0; released as 05_validated_stats.json), the lowest count being 24. The check therefore excluded no case, and the analytic cohort follows from the pinned files alone. We retain the count as the per-case descriptor pmc_article_count (Section 4).

For each case we assembled a 50-gene candidate list: the causal gene plus 49 distractors drawn uniformly at random, without replacement, from the pinned HGNC protein-coding set (19,296 symbols, HGNC snapshot 2026-04-07; [26]), excluding the causal gene. We fixed the list at 50 on three grounds: a filtered exome typically leaves of the order of 10 to 100 protein-altering candidates per case after standard quality, frequency and American College of Medical Genetics and Genomics (ACMG) guideline filtering [27,28]; rare Mendelian disease is usually caused by variants in a single protein-coding gene [29], and the cohort admits only single-gene phenopackets, so a single-target list is faithful to the ground truth; and listwise ranking quality degrades for longer candidate lists [30]. A fixed length also keeps rank-based metrics comparable across cases. This controlled setting is an evaluation benchmark, not a genome-wide clinical simulation (Section 5).

Sampling uses a per-case derived seed: we read the 8-byte BLAKE2b digest of the ASCII string “42|{case_id}” as a big-endian unsigned integer and seed a Python random.Random instance with it. Any single case therefore regenerates independently and bit-identically, without resampling the others. We draw the 49 distractors with random.sample from the pool sorted by symbol (sorting is required for a byte-stable draw), prepend the causal gene, and shuffle the 50-gene list with the same generator. We record the causal gene’s resulting position as causal_gene_index_in_candidates. Three sampled neurological cases have a small nuclear RNA gene as their causal gene (two *RNU4-2*, one *RNU2-2*), which falls outside the protein-coding pool; we therefore dropped them at this stage. The sample reduces from 1,050 to the final *n* = 1,047 (250 developmental, 300 immunological, 250 metabolic, 247 neurological). We call this the standard candidate-list variant: random distractors from the full protein-coding space are a standard genome-wide design.

#### Hard candidate-list variant

To provide a difficulty axis orthogonal to the leakage axis, we additionally release a hard variant in which the 49 distractors are the genes *phenotypically most similar* to each case rather than random. For every HGNC protein-coding gene with HPO annotations, we score phenotypic similarity to the case as the best-match-average of Resnik term similarities, where Resnik similarity between two HPO terms is the information content of their most informative common ancestor [31]. We aggregate the term-level scores symmetrically. The forward score averages, over the case’s HPO terms, each term’s best Resnik match among the gene’s annotated phenotypes (genes_to_phenotype, HPO v2026-02-16). The reverse score averages, over the gene’s annotated terms, each term’s best match among the case’s terms. We average the two with equal weight. Information content is IC(*t*) = − ln(*n_t_*/*N*), where *N* is the number of HPO-annotated genes and *n_t_* the number whose annotations, propagated over the is_a closure of the ontology, include *t*; obsolete terms are mapped to their primary identifiers via alt_id, and terms with IC = 0 contribute nothing. We score only genes sharing at least one IC *>* 0 ancestor with the case; genes sharing none score zero by construction, and we discard them. We take the top-49 genes by this score as distractors, excluding the causal gene and any gene annotated to the case’s own causal disease(s), so that a distractor can never be a genuine alternative cause. We break ties by gene symbol and shuffle the final 50-gene list with the *same* per-case BLAKE2b seed as the standard variant. The hard variant is therefore equally deterministic and case-paired: case_id, causal_gene, hpo_terms and the metadata layers are identical, and only candidate_genes differ. All 1,047 cases yielded *≥* 49 scored candidates, so no random fill was required.

The two variants are archived as sibling datasets (Table 3; Data Citations 1 and 2); their difficulty is contrasted in Section 4.

**Table 3:**
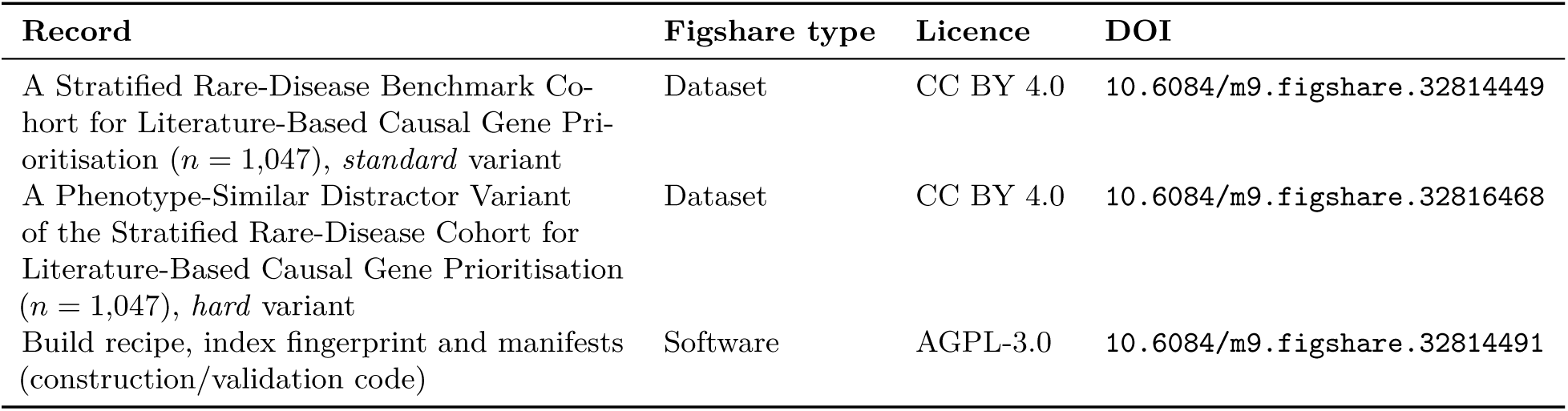
Archived records and persistent identifiers for the released resource.

| Record | Figshare type | Licence | DOI |
| --- | --- | --- | --- |
| A Stratified Rare-Disease Benchmark Cohort for Literature-Based Causal Gene Prioritisation ($n = 1,047$), *standard* variant | Dataset | CC BY 4.0 | 10.6084/m9.figshare.32814449 |
| A Phenotype-Similar Distractor Variant of the Stratified Rare-Disease Cohort for Literature-Based Causal Gene Prioritisation ($n = 1,047$), *hard* variant | Dataset | CC BY 4.0 | 10.6084/m9.figshare.32816468 |
| Build recipe, index fingerprint and manifests (construction/validation code) | Software | AGPL-3.0 | 10.6084/m9.figshare.32814491 |

### 2.2 Stratification metadata

#### Annotation-overlap flag

The goal of this flag is to mark, for each case, whether a tool curated from the HPO annotations could have seen that case’s source publication. For each case we computed a binary annotation_overlap flag. The flag is 1 if the case’s source PMID appears in phenotype.hpoa v2026-02-16 as a reference for any annotation of any of the case’s causal OMIM disease IDs, and 0 otherwise. We parse the PMID from the case_id, whose Phenopacket Store convention encodes the source publication as a PMID_<digits>_… filename stem. We parse the phenotype.hpoa rows into unique (disease_id, reference) keys, splitting multi-reference cells on the semicolon separator and retaining only PMID:-prefixed references, then join each case against this index. All 1,047 cases resolved to both a PMID and an OMIM disease ID, so no case fell into the overlap-absent subset merely for want of a joinable identifier. Because we compute the flag from a case’s source PMID and disease identifiers, cases that share a source publication and disease necessarily share a flag value. The 1,047 cases derive from 415 unique PMIDs, so the flag varies across substantially fewer independent units than cases.

The flag partitions the cohort into an overlap-present subset (*n* = 765; 73.1 %) and an overlap-absent subset (*n* = 282; 26.9 %), for which we detect no source-publication overlap with the annotation file. The overlap-absent subset supports a fairer comparison against curated tools, but does not guarantee the absence of leakage; we set out the residual routes under Limitations (Section 5).

#### Publication-recency strata

We retrieved source-publication dates for the 415 unique cohort PMIDs from NCBI E-utilities (efetch, PubMedPubDate with PubStatus=“pubmed”). All 415 resolved (100 %; oldest 1988, most recent 2025, median 2018 over unique PMIDs). We split cases at 2020-01-01 into pre-2020 (*n* = 601) and post-2020 (*n* = 446) strata. We provide the crossed post-2020 *×* overlap-absent subset (*n* = 88) as the closest available approximation to a “novel-association” cohort.

### 2.3 PMC Open Access corpus acquisition and filtering

We built the retrieval substrate from the PMC OA full-text XML corpus (the open, redistributable portion of PubMed Central; [32]), taking all three licence tiers of the NCBI FTP bulk distribution (oa_comm, oa_noncomm, oa_other): the 2026-01-23 baseline packages plus every incremental package up to the 2026-05 snapshot date.

Article licences differ both across and within these tiers (CC BY, CC BY-NC, and other terms), so we do not redistribute the verbatim chunk text (Section 3, Section 5). We provide the build recipe and fingerprints instead. The three steps below are implemented in scripts/corpus/02_extract_ and_parse_ftp.py and scripts/corpus/03_normalize_dedupe_filter.py.

#### Parsing and retraction exclusion

We parsed each tarball from JATS XML into one record per article, retaining identifiers, bibliographic metadata, JATS subject categories, abstract, and section-segmented body text. We excluded retracted articles *before* parsing using the Retracted column of the .filelist.csv manifest that NCBI ships alongside each tarball: we dropped any article whose AccessionID is flagged Retracted=yes and wrote the dropped identifiers to an audit log (parsed/skipped_retractions.jsonl). We also dropped articles whose XML failed to parse and articles carrying no PMC identifier.

#### Deduplication

Baseline and incremental packages both contain an article whenever it has been updated, so we deduplicated records by PMC identifier with the most recent package taking precedence: we visit input files in reverse chronological order (newest incremental first, baselines last) and emit each PMC identifier at most once. Deduplication therefore depends only on the pinned snapshot, not on processing order within it.

#### Genetics-relevance filter

We decided relevance by lexical matching rather than ontology lookup: PMC OA JATS packages do not carry MeSH indexing, so no MeSH descriptors were available at this stage. For each article we formed a single case-folded haystack from three fields—the article title, the abstract, and the JATS <subject> category strings. We did *not* use body text. We retained an article if either (a) the haystack contains, as a substring, any term from a pinned 77-term genetics vocabulary (covering general genetics and genomics terms such as *genetic*, *genome*, *mutation*, *variant*, *allele*, *exome*, *phenotype*; inheritance and variant-class terms such as *autosomal*, *X-linked*, *de novo*, *missense*, *frameshift*, *splice*; rare-disease terms such as *rare disease*, *orphan disease*, *Mendelian*, *monogenic*, and a set of named Mendelian conditions; and assay and regulation terms such as *RNA-seq*, *methylation*, *chromatin*); or (b) the haystack matches a word-boundary regular expression covering gene-structure nouns (*gene*, *locus*, *exon*, *intron*, *codon*, *protein*, *amino acid*), sequencing and editing technologies (*sequencing*, *Sanger*, *Illumina*, *Nanopore*, *CRISPR*, *Cas9*, *TALEN*, *ZFN*), numeric database identifiers (OMIM:, HGNC:, Orphanet:, MONDO:, HPO: followed by digits), cytogenetic locations (e.g. chr17p13), and sequence-length units (bp, kb, Mb). We dropped articles with an empty haystack. The exact vocabulary and regular expression are the literals GENETICS_VOCAB and GENETICS_REGEX in scripts/corpus/03_normalize_dedupe_filter.py, as archived in the code deposit (Data Citation 3, release tag paper-methods-v1.4); they are the normative definition of the filter, and the prose above is a summary of them.

Several vocabulary terms are general biomedical words, and matching is by substring rather than by word boundary. The retained set therefore includes literature outside rare-disease genetics, such as somatic-variant and viral-variant papers, and topical precision is determined by the query-time ranker rather than by the filter.

#### Corpus funnel

Applying the pipeline to the pinned snapshot yielded 7,870,943 parsed article records. Deduplication left 7,809,296 unique PMC identifiers, dropping 61,471 duplicate and 176 no-identifier records. The relevance filter then removed 5,554,908 of these, leaving 2,254,388 articles (*∼*2.25 million) as the indexed corpus.

We record these counts in parsed/_normalize_stats.json and release the full list of retained PMC identifiers as retained_pmcids.txt (2,254,388 lines, one PMCxxxxxxx identifier per indexed article). Readers can therefore audit the relevance filter and verify a rebuild against the exact indexed set without rebuilding the index.

### 2.4 Hybrid retrieval index construction

The deterministic build pipeline is summarised in Figure 2 and is implemented in scripts/corpus/ 04_chunk_normalized.py. Chunking is *section-aware*: we treat an article’s abstract as its first section and each body section as a subsequent one, and no chunk spans a section boundary, so every chunk carries an unambiguous section label. We skip sections with fewer than 50 characters of text. We emit a section of at most 512 tokens as a single chunk containing the section text verbatim. We split a longer section with a sliding window over its token sequence, using a window of 512 tokens and a stride of 462 (a 50-token overlap), and decode each window back to text. Chunks of long sections are therefore tokeniser round-trips rather than verbatim substrings. Tokenisation uses the PubMedBERT tokeniser [33].

**Figure 2:**
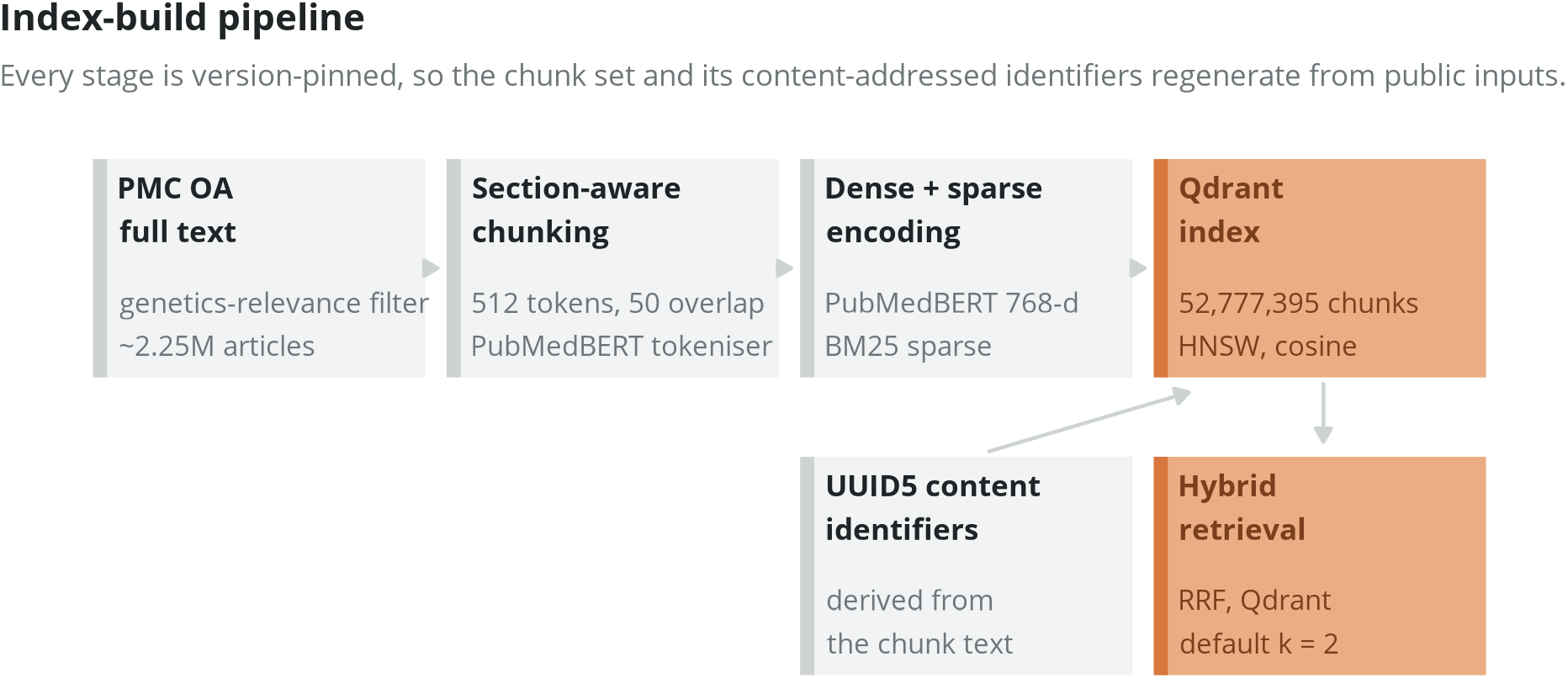
Deterministic index-build pipeline. A genetics-relevant subset of the PMC Open Access corpus (selected by a lexical genetics-relevance filter over title, abstract and JATS subject categories) is parsed from full-text XML, split into 512-token chunks with 50-token overlap using the PubMedBERT tokeniser, and assigned content-addressed UUID5 identifiers. Each chunk receives a PubMedBERT dense embedding and a BM25 sparse representation; both are stored in a Qdrant collection (52,777,395 chunks). At query time, dense and sparse hits are combined by Reciprocal Rank Fusion, with an optional MedCPT cross-encoder reranker. Every stage is version-pinned so the same chunk set and identifiers regenerate from the same public inputs.

Each chunk receives a content-addressed UUID5 identifier, computed as

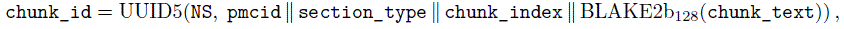

where the four fields are joined by the | character, BLAKE2b_128_ is a 16-byte BLAKE2b digest rendered as lowercase hexadecimal, and NS is the pinned namespace UUID 6f9619ff-8b86-d011-b42d-00cf4fc964ff. The namespace is fixed for the lifetime of the collection: changing it would invalidate every existing point identifier. Because identifiers depend only on the chunk’s content and position and not on processing order, re-running the pipeline over the same pinned snapshot reproduces the same identifiers, and upserts into Qdrant are idempotent.

We store two complementary representations for every chunk. The *dense* representation is a 768-dimensional PubMedBERT embedding [33]—a biomedical model in the BERT family [34,35]— mean-pooled and L2-normalised, computed in half precision on GPU. The *sparse* representation is a BM25 vector computed with fastembed’s Qdrant/bm25 model. Both live in Qdrant v1.14.1 [36]. At query time, Qdrant’s built-in Reciprocal Rank Fusion (RRF) merges the two rankings without tuning [37]: a document *d* scores Σ*_i_* 1*/*(*k* + *r_i_*(*d*)), where *r_i_*(*d*) is its zero-based rank in the dense and the sparse result list. On the pinned v1.14.1 the engine constant is *k* = 2 and is not configurable; a settable *k* arrived only in Qdrant v1.16.0. We report the engine default rather than a tuned constant, which is consistent with this resource’s aim of characterising an untuned substrate.

Document-side sparse vectors carry term frequencies only. Qdrant applies the inverse-document-frequency weighting server-side with Modifier.IDF from collection statistics, so queries use the term-frequency-only encoder. The production collection (geno_agent_pmc_oa_v1) contains 52,777,395 chunks. We use this value as the index fingerprint, verified via the Qdrant points_count API and recorded in data/MANIFEST.tsv. The concrete index parameters needed to reproduce the substrate at the configuration level are given in Table 5. A MedCPT cross-encoder [38] is provided as an optional query-time reranker; it is part of the retrieval substrate but is not required to reproduce the index.

## 3 Data Records

The resource is openly archived on Figshare with persistent DOIs (Table 3): the standard-variant cohort (Data Citation 1), the phenotype-similar hard variant (Data Citation 2), and the build recipe with the index fingerprint and validation records (Data Citation 3).

### Cohort dataset contents

The canonical file test_cases.jsonl holds one JSON object per case with the fields in Table 4; the metadata sidecars (annotation_overlap.json, pmid_dates.json), the staged provenance files (01_all_phenopackets.jsonl through 06_with_candidates.jsonl), a build manifest with the SHA-256 and byte size of test_cases.jsonl, and per-file checksums are included. A Croissant (MLCommons) machine-readable dataset description (croissant.json) is included in each deposit for automated discovery and ingestion.

**Table 4:**
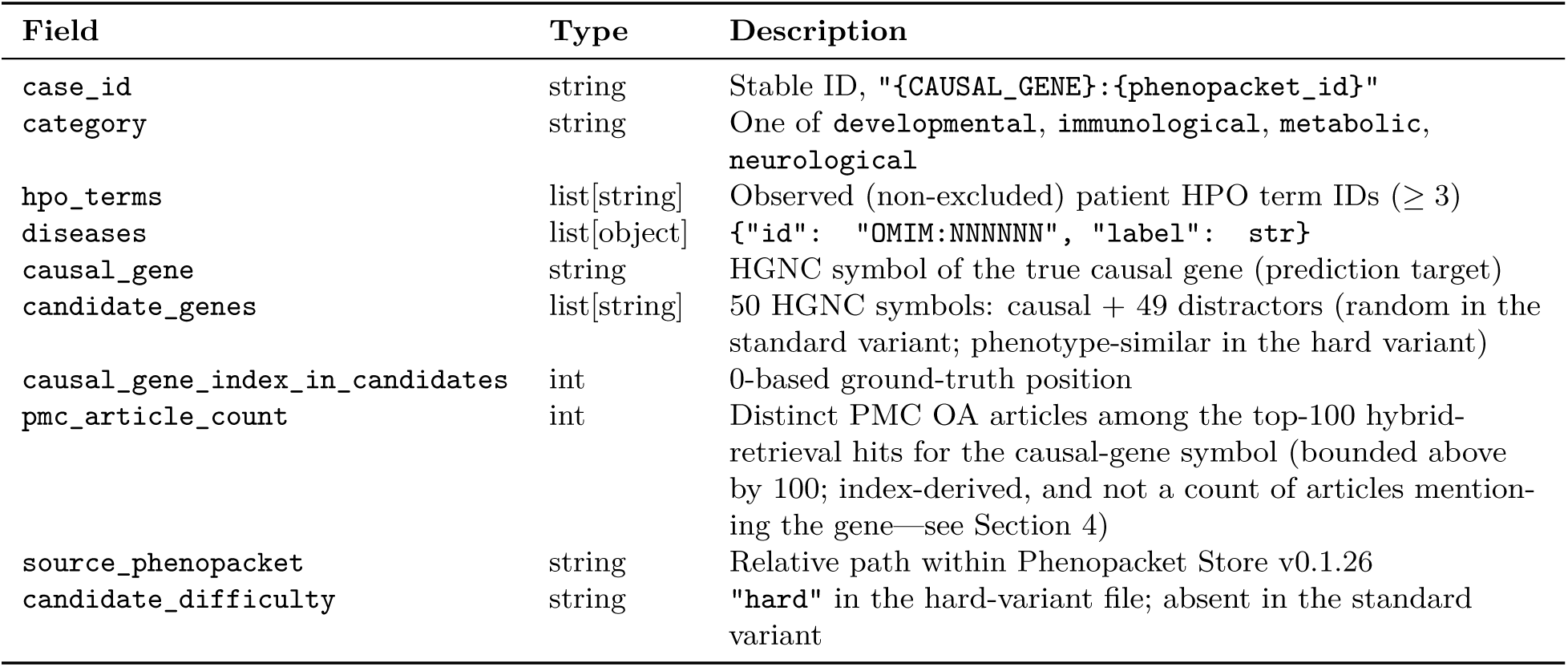
Per-case schema of test_cases.jsonl.

The hard-variant file (test_cases_hard.jsonl) shares every field above with the standard cohort except candidate_genes and causal_gene_index_in_candidates, and adds candidate_ difficulty; a per-case selection-diagnostics sidecar (hard_candidates_stats.json) records the causal– and distractor-similarity scores. The hard variant ships its own build manifest (test_ cases_hard_manifest.json) recording the SHA-256 and byte size of (test_cases_hard.jsonl), following the same convention as the standard variant.

### Retrieval index

The 323 GB Qdrant index and the verbatim PMC OA chunk text are recipe-only (mixed-licence source text): the methods record provides the deterministic build pipeline (Figure 2) and the fingerprint (52,777,395 chunks; SHA-256 of upstream inputs in data/MANIFEST.tsv) so that the index regenerates rather than being hosted. The configuration parameters a reuser needs are in Table 5 and the pinned upstream versions in Table 6. For backward compatibility with existing point payloads, the Qdrant payload field named mesh_terms contains JATS <subject> category strings, not MeSH descriptors; PMC OA JATS packages do not carry MeSH indexing.

**Table 5:**
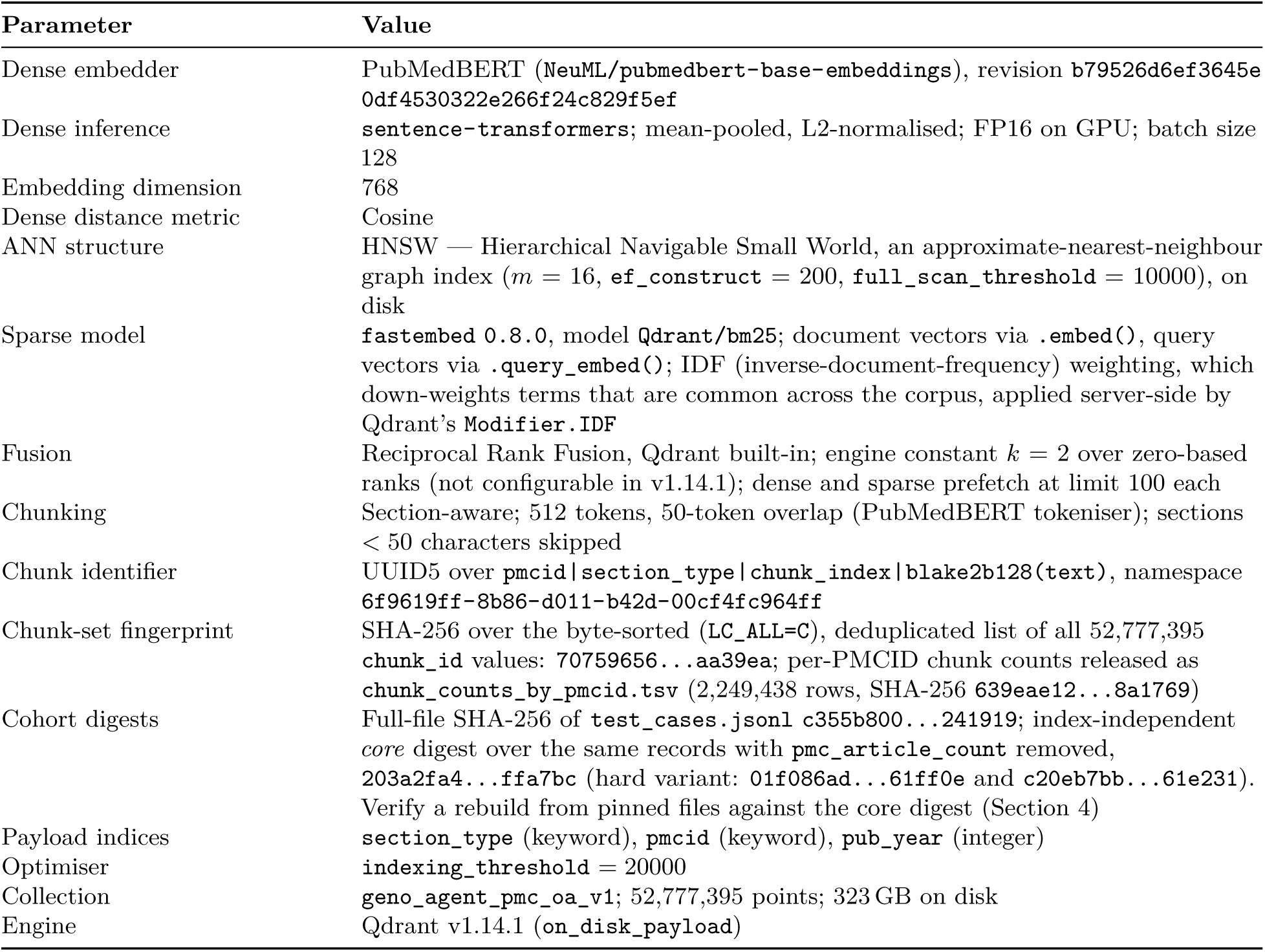
Retrieval-index configuration and released digests (reproduction parameters).

**Table 6:** Pinned upstream inputs (provenance).

| Input | Pinned version | Licence |
| --- | --- | --- |
| GA4GH Phenopacket Store | v0.1.26 (2026-01-13) | CC BY 4.0 |
| Human Phenotype Ontology | v2026-02-16 | open (HPO) |
| Mondo Disease Ontology | v2026-03-03 | CC BY 4.0 |
| HGNC complete set | 2026-04-07 | open (EBI/HGNC) |
| PubMedBERT embedder | NeuML/pubmedbert-base-embeddings, rev. b79526d6 | open |
| fastembed (BM25 sparse) | 0.8.0, model Qdrant/bm25 | Apache 2.0 |
| Qdrant | v1.14.1 | Apache 2.0 |
| PMC Open Access subset | retrieved 2026-05 | mixed (per-article) |

## Data availability

The cohort files are openly available under CC BY 4.0 from Figshare, with no restrictions on access or reuse: the standard variant at https://doi.org/10.6084/m9.figshare.32814449 (Data Citation 1) and the phenotype-similar hard variant at https://doi.org/10.6084/m9.figshare.32816468 (Data Citation 2). The build recipe, manifests and validation records are available under AGPL-3.0 at https://doi.org/10.6084/m9.figshare.32814491 (Data Citation 3). The verbatim PMC Open Access chunk text and the Qdrant index are the one exception, for the licensing reason given above; both regenerate from the released recipe and the public inputs of Table 6.

## 4 Technical Validation

### Eligibility and determinism

Every retained case satisfies the inclusion criteria above, all of which are evaluated against pinned files alone. A single index-dependent coverage check ran at the sampling stage and excluded no case (Section 2.1), so the cohort as released does not depend on the state of the retrieval index. The screening funnel (9,588 phenopackets loaded *→* 6,382 passing inclusion criteria (i)–(ii) and the disease-scope exclusions *→* 4,670 categorised *→* 1,050 stratified sample *→* 1,047 final, after removing three non-protein-coding causal genes) and the final per-category counts (250/300/250/247) are regenerable from the pinned inputs and seed (Figure 1; Table 7).

**Table 7:** Cohort descriptors by disease category (all values regenerable from the released files). “Overlap” columns give the annotation-overlap split; “Recency” columns the pre/post-2020 source-publication split; medians are per case. “Med. retr. articles” is the median pmc_article_count, i.e. the number of distinct articles among the top-100 hybrid-retrieval hits for the causal-gene symbol (bounded above by 100), not a count of articles mentioning the gene; the cohort minimum is 24, so every case clears the five-article check applied at sampling (Section 2.1). Cases outnumber source publications (1,047 cases from 415 unique PMIDs), so cases are clustered within publications; see Usage Notes. The per-category unique-PMID counts sum to 416 rather than 415 because one publication (PMID 37964426) contributes cases assigned to two categories.

| Category | *n* | Unique<br>PMIDs | Overlap |  | Pre-2020 | Post-2020 | Med. HPO<br>terms | Med. retr.<br>articles |
| --- | --- | --- | --- | --- | --- | --- | --- | --- |
|  |  |  | present | absent |  |  |  |  |
| Developmental | 250 | 91 | 158 | 92 | 161 | 89 | 9 | 55 |
| Immunological | 300 | 107 | 259 | 41 | 180 | 120 | 8 | 52 |
| Metabolic | 250 | 85 | 160 | 90 | 160 | 90 | 7 | 62 |
| Neurological | 247 | 133 | 188 | 59 | 100 | 147 | 8 | 55 |
| **Total** | **1,047** | **415** | **765** | **282** | **601** | **446** | **8** | **55** |

The released descriptor pmc_article_count requires a caveat. It counts distinct articles within a fixed-size (top-100) hybrid-retrieval result set, so it is bounded above by 100, does not distinguish a gene with hundreds of publications from one with thousands, and does not require the gene symbol to appear in the retrieved text. Every case clears the five-article check applied at sampling by a wide margin (cohort minimum 24, median 55; Table 7). Read the descriptor as a measure of retrieval-result diversity rather than of per-gene corpus coverage; users needing the latter should compute it from the released per-PMCID chunk manifest (Section 3).

### Cohort characterisation

Figure 3 summarises the cohort. The per-category counts follow the sampling design, not the composition of the eligible pool (Figure 3a). The annotation-overlap analysis shows that **73.1** % of cases (765*/*1,047) are overlap-present (Figure 3b)—direct evidence that annotation overlap is the common case, not a corner case, and motivating the overlap-absent subset. The source publications span 1988–2025 (median 2018 over the 415 unique PMIDs; 601 pre-2020 / 446 post-2020 cases; Figure 3c), and phenotype-annotation depth has a median of 8 HPO terms per case (range 3–43; Figure 3d). All descriptors are reproducible from annotation_overlap.json, pmid_dates.json and the category/hpo_terms fields.

**Figure 3:**
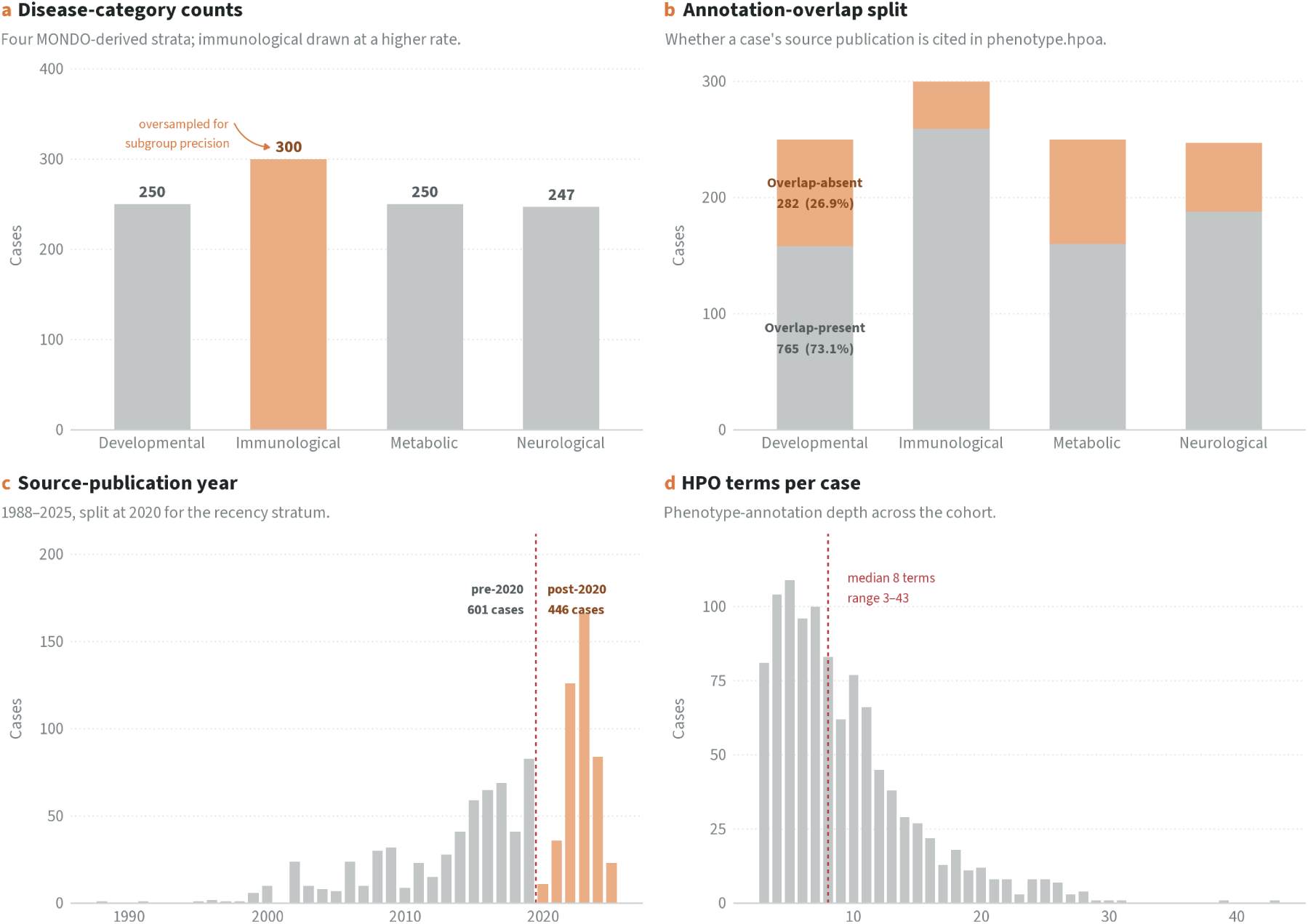
Cohort characterisation. (a) Cases per disease category (250*/*300*/*250*/*247). (b) Annotation-overlap split per category—overlap-present (leakage risk) vs. overlap-absent (*n* = 282). (c) Source-publication year distribution with the 2020 stratification boundary. (d) Distribution of HPO terms per case (phenotype-annotation depth).

### Candidate-list difficulty (standard vs hard)

The two candidate-list variants realise a controlled difficulty axis, quantified on a common scale—the case-to-gene Resnik best-match-average (BMA) phenotypic similarity (Figure 4). In the *standard* variant, random distractors are mostly phenotypically unrelated to the case (median distractor BMA 0.00 vs. a median causal-gene BMA of 2.36; Figure 4a), so the causal gene is usually the most concordant candidate; even so the task is not trivial, as at least one random distractor matches or exceeds the causal gene’s phenotypic similarity in **7.4** % of cases (Figure 4b). Because a tie could in principle be degenerate—both the causal gene and the hardest distractor scoring 0.00 where the causal gene carries no informative HPO annotation—we report the decomposition: all 78 standard-variant cases (7.45 %) are strict exceedances, with no ties at zero and none above zero. Exactly one cohort case has a causal-gene BMA of 0.00, and there too a distractor scores strictly higher. The hard variant behaves the same way (448 cases, 42.79 %, all strict). The split is released as difficulty_tie_split.json. The *hard* variant raises this sharply: its distractors are strongly phenotype-similar (median distractor BMA 1.71), and a distractor matches or exceeds the causal gene in **42.8** % of cases—a roughly six-fold increase in genuinely confusable candidate lists. The same-disease exclusion removes any gene annotated to the case’s own disease(s), so no distractor is a curated alternative cause for that diagnosis. We treat distractors as negatives relative to the single recorded causal gene; for cases with digenic or oligogenic contributions that labelling may be incomplete (see Limitations). Because the variants share cases, seed and ground truth, the difference is attributable to distractor selection alone.

**Figure 4:**
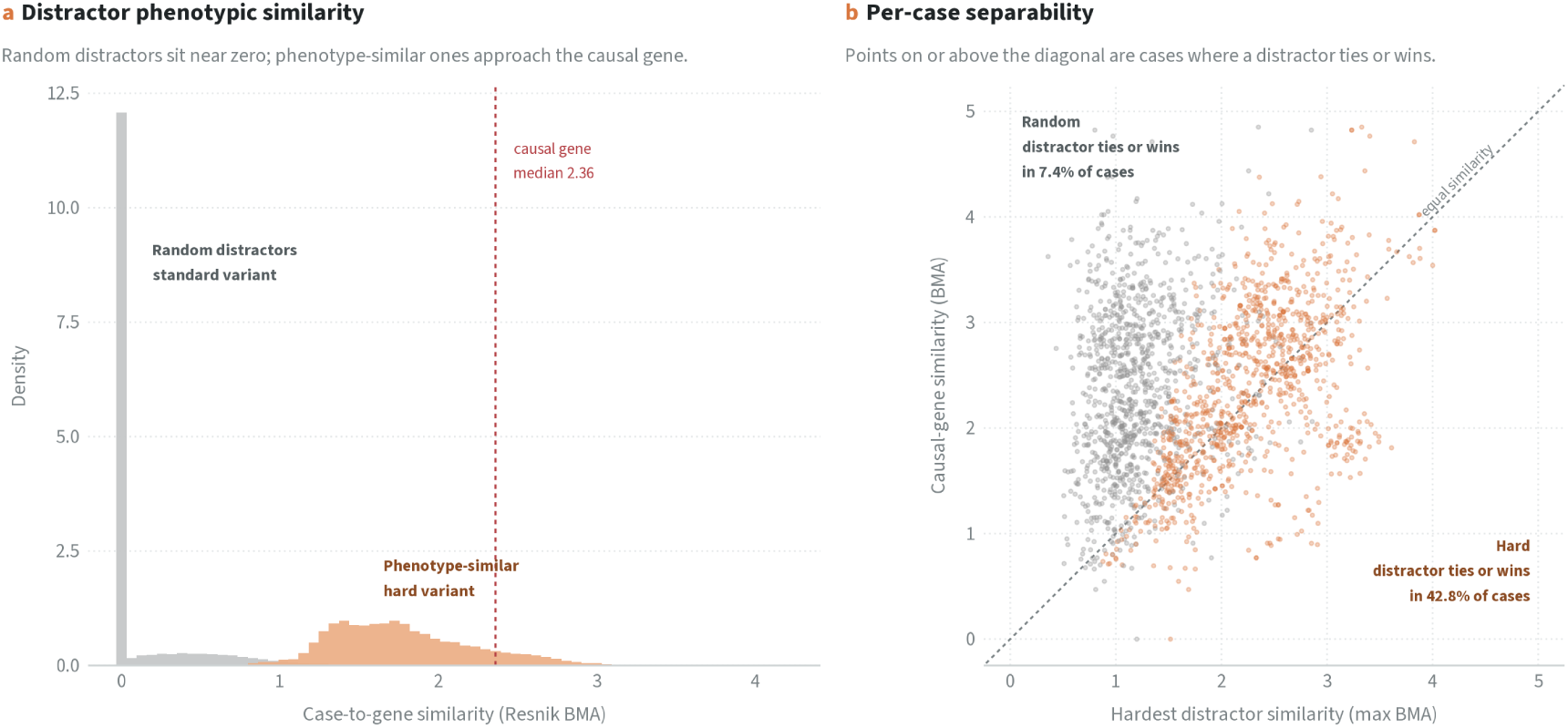
Candidate-list difficulty: standard (random) vs hard (phenotype-similar) distractors, on a common Resnik best-match-average (BMA) phenotypic-similarity scale. (a) Distractor-similarity distributions—random distractors (grey) cluster near zero, hard distractors (orange) are shifted toward the causal-gene median (dashed red). (b) Per-case causal-gene similarity vs. the hardest distractor; points on/above the diagonal are cases where a distractor ties or exceeds the causal gene (7.4 % standard vs. 42.8 % hard).

### Reproducibility of the resource

Determinism of the released resource rests on (i) explicit sorting at every step where iteration order could matter, together with BLAKE2b-derived identifiers in place of Python’s salted hash; (ii) UUID5 content-addressed chunk identifiers; (iii) seed-42 sampling at every stochastic step, including the per-case BLAKE2b-derived distractor seed, so that individual cases regenerate independently; and (iv) fully pinned dependency versions. We export PYTHONHASHSEED=42 for child processes, but no released artefact depends on it. Every cohort-construction stage—phenopacket ingestion, inclusion, MONDO categorisation, seeded stratified sampling, candidate-list construction and finalisation—depends only on pinned files and seeds.

Regenerating the cohort reproduces the SHA-256 recorded in each build manifest (standard c355b800…241919; hard 01f086ad…61ff0e). Those digests cover the full canonical file, which includes the index-derived pmc_article_count, and a rebuild on different hardware may return slightly different values for that descriptor. We therefore also release a *core* digest over the same records with the field removed (203a2fa4…ffa7bc; hard c20eb7bb…61e231). It is the digest that certifies the benchmark, since the descriptor enters no metric and no inclusion decision: reusers working from the pinned files alone should verify against it, while the full-file digest reproduces only against the same index.

For the retrieval index we make a narrower claim. Chunking is deterministic and chunk identifiers are content-addressed, so re-running the pipeline over the same pinned PMC OA snapshot yields the same chunk set under the same identifiers. We verify this at two levels: the collection point count matches the recorded 52,777,395, and the SHA-256 over the byte-sorted, deduplicated identifier list (70759656…aa39ea) fingerprints the chunk set itself, since any different chunk boundary, retained article or section segmentation would change it. Deduplication is what makes that digest a property of the chunk set rather than of a particular run: a resumed build can re-emit a record an earlier pass already wrote, and it upserts onto the same point. Our own build emitted 52,782,789 records, of which 5,394 were exact identifier duplicates, leaving 52,777,395 distinct identifiers—precisely the recorded point count, which independently confirms that the upserts were idempotent. The accompanying per-PMCID manifest (chunk_counts_by_pmcid.tsv, 2,249,438 rows, SHA-256 639eae12…8a1769) localises any discrepancy to specific articles without redistributing chunk text; it covers fewer identifiers than the 2,254,388 retained articles because 4,950 of them have no section reaching the 50-character minimum and so contribute no chunk.

The claim stops at content. HNSW graph construction depends on insertion order and the build uses parallel, resumable upserts, so two builds from identical inputs are content-equivalent rather than byte-identical on disk; the stored vectors likewise differ in their last bits across environments, because dense embeddings were computed in half precision (FP16) on GPU. What regenerates and is verifiable is which chunks exist, under which identifiers, and with which pinned model and parameters. These checks establish reproducibility of the resource itself; the end-to-end robustness of any system built on it is a property of that system.

### Retrieval-substrate validation

The checks above establish that the indexed content is regenerable, not that it is *useful*. We therefore report two index-level characterisations (Table 8), both properties of the corpus and the retrieval configuration alone: no ranking model, language model or prioritisation tool is involved. The binding constraint is corpus coverage rather than ranking: of the 415 unique source publications behind the cohort, 174 have no PMC record at all and a further 111 fall outside the PMC OA subset or the genetics-relevance filter, leaving 130 (31.3 %) in the index and 345 of the 1,047 cases eligible for Check A. Among those eligible cases a single unrefined query—a bare causal-gene symbol, or the case’s HPO term labels joined by spaces—recovers the source article among the parent articles of the top-100 chunks for 35.7 % and 37.4 % of cases respectively; the phenotype-label query leads the gene-symbol query at every cut-off. Check B finds that a mean of 18.6 % (median 18.0 %) of the top-100 chunks returned for a gene symbol contain that symbol under a case-sensitive word-boundary match, ranging from 2 % to 50 % across the 100 sampled genes, with no gene returning zero literal matches—the substrate is lexically grounded for every gene tested, while the majority of hybrid hits are semantically rather than literally related. Both queries are unrefined, so Check A is a lower bound on retrievability rather than a ceiling. Most source publications are absent from the open-access corpus, so a literature-based tool evaluated on this cohort will usually not be able to recover the answer by locating the originating report. These values characterise the substrate, not any system built on it.

**Table 8:** Index-level retrieval-substrate characterisation. Both checks use the hybrid configuration of Table 5 (dense PubMedBERT and BM25 sparse prefetch, each at limit 100, fused by Qdrant’s built-in Reciprocal Rank Fusion with the engine constant *k* = 2), identical to the configuration used to compute pmc_article_count. Recall is over the 345 cases whose source article is present in the index; the remaining cases are not scored, since a source article absent from the corpus cannot be retrieved from it.

| Check | Query | Metric | Value |
| --- | --- | --- | --- |
| Source articles present in index | — | $n$ / 415 unique source PMIDs | 130 (31.3%) |
| A. Source-article recall | causal-gene symbol | recall@10 / @50 / @100 | 0.171 / 0.278 / 0.357 |
| A. Source-article recall | HPO term labels | recall@10 / @50 / @100 | 0.229 / 0.310 / 0.374 |
| B. Symbol grounding | causal-gene symbol | mean / median fraction of top-100 chunks containing the symbol (100 genes, seed 42) | 0.186 / 0.180 |

### Independence of the annotation-overlap layer

The annotation_overlap flag depends only on public inputs (the case PMID and phenotype.hpoa) and not on any tool’s output, so it is a property of the benchmark rather than of a particular system, and can be recomputed by any user against a different phenotype.hpoa release.

## 5 Usage Notes

### Recommended evaluation protocol

For fairer comparison of a literature-based tool against curated tools (e.g. those drawing on phenotype.hpoa), report metrics on the overlap-absent subset (*n* = 282) as the primary endpoint, with the full cohort as a supportive secondary analysis. Top-*k* accuracy, Mean Reciprocal Rank and NDCG@10 over the 50-gene candidate list are the natural metrics; we release the candidate list and the ground-truth index so that any ranking system can be scored identically. Note that with exactly one relevant item per case, NDCG@10 is a monotone transform of the reciprocal rank truncated at 10; we retain it for comparability with information-retrieval conventions rather than as an independent view of performance. The flag is computed against phenotype.hpoa v2026-02-16 and is release-specific: as the annotation file grows, more source publications become cited and the overlap-absent subset shrinks. Users should recompute the flag against the annotation release contemporaneous with the tool under evaluation; the procedure requires only public inputs (Section 2.2).

### Clustered cases

The 1,047 cases derive from 415 unique source publications—a median of one case per publication, but a mean of 2.5 and a maximum of 42, so the distribution is strongly right-skewed. Cases from the same publication are not independent: they share a source, frequently a causal gene, and by construction an annotation-overlap flag. Per-case metrics treated as independent observations will therefore have understated variance, and the effective sample size for any leakage-stratified comparison is closer to the number of publications than of cases. The effect is sharpest in the smallest recommended cell: the crossed post-2020 *×* overlap-absent subset contains 88 cases but only 18 unique publications, one contributing 35 of the 88, so read it as a qualitative probe rather than a powered comparison. We recommend clustering confidence intervals and significance tests on source PMID—for example a publication-level bootstrap that resamples publications with replacement and retains all their cases. The case_id field encodes the source PMID, so the clustering variable requires no additional file; per-stratum unique-publication counts are released in clustering_stats.json.

### Weighting

The four strata were sampled at very different rates (Table 2): from 77 % of the eligible immunological pool down to 8 % of the neurological pool. Cohort-level pooled metrics therefore reflect the sampling design, not the eligible population. Report per-category metrics as the primary per-domain result; where a single cohort-level number is wanted, compute it as a stratum-weighted estimate using the released inclusion probabilities. The same applies within the overlap-absent subset, whose category composition (92*/*41*/*90*/*59) departs further still from the drawn proportions and whose immunological stratum falls to *n* = 41. Per-category estimates there are underpowered for immunological cases; report that subset as a whole, or as a stratum-weighted estimate, rather than as four per-category numbers.

### Recency analysis

Use the publication-year split (or the crossed post-2020 *×* overlap-absent subset, *n* = 88) to probe generalisation to associations that post-date curation cycles. The crossed cell is the smallest and least balanced unit in the resource and should be sized before it is used: its category composition is 20*/*7*/*23*/*38, so no per-category estimate is supportable within it, and it is additionally the most strongly clustered stratum (see *Clustered cases* above).

### Difficulty axis (standard vs hard)

Because the standard and hard variants share cases, ground truth, seed and metadata layers, they form a difficulty axis orthogonal to the leakage axis, supporting a 2 *×* 2 (difficulty *×* leakage) evaluation: each system can be scored on the standard and hard candidate lists, each split by the overlap-absent subset. The standard variant measures genome-wide discrimination; the hard variant stresses differential-diagnosis behaviour against phenotype-similar competitors—subject to the tool-class asymmetry noted under Limitations. We recommend reporting both, with the hard *×* overlap-absent cell as the most stringent condition.

### Rebuilding the index

We do not distribute the 323 GB index. The underlying articles carry mixed licences that do not permit redistributing the verbatim chunk text, and a static download could not be verified or updated by reusers. We release instead the scripts, the pinned upstream versions (Table 6), the configuration (Table 5) and the fingerprints needed to rebuild it (52,777,395 chunks plus SHA-256 manifests of the upstream inputs). A reuser can regenerate the chunk set from public sources, check it against the released digest, and re-run the build against a newer PMC OA snapshot. Such a rebuild reproduces the chunk set under the same identifiers; it does not reproduce a byte-identical index or identical vectors (Section 4). Because PMC OA grows over time, users targeting an identical chunk set should pin the same snapshot date.

Verify a rebuild by recomputing the sorted chunk-identifier digest under LC_ALL=C and comparing against the released value; chunk_counts_by_pmcid.tsv localises any mismatch to specific PMC identifiers. Both the byte-ordering locale and the deduplication step are load-bearing: a digest computed under a different collation, or over the raw record stream rather than the distinct identifier set, will not match.

### Licensing

The cohort is CC BY 4.0 (it derives from the CC BY 4.0 Phenopacket Store and open ontologies). The build/evaluation code is AGPL-3.0. Verbatim PMC OA chunk text is not redistributed because article licences are mixed; users regenerate it from PMC under the source licences.

### Scope

This resource describes phenotype-driven prioritisation inputs (HPO terms + candidate lists). It does not include patient variant calls; variant-aware benchmarking is out of scope.

### Limitations

Six limitations bound the claims a user should draw from this resource. First, the single-causal-gene, fixed-50 design is an evaluation harness rather than a model of clinical diagnosis (Section 2.1). Each case labels exactly one causal gene and 49 distractors, yet some cases may have more than one plausible or contributing gene—digenic or oligogenic contributions, or phenotypically overlapping differential diagnoses—and 50 candidates do not represent the genome-wide space a tool faces in practice. The choice of 50 rather than, say, 100 is pragmatic, and results may shift modestly with list length. Users who need genome-scale ranking or multi-gene ground truth should treat these lists as a controlled stress test.

Second, the overlap-absent subset is a conservative proxy for a leakage-free set, and no more than that. The annotation_overlap flag detects one specific, checkable event: that a case’s source PMID is cited in phenotype.hpoa for the case’s own disease(s). Several routes to prior exposure escape it—the same case or family curated under a related or renamed disease entry; an annotation curated under an ORPHA or DECIPHER entry for the same condition, since the join resolves OMIM identifiers only; a later publication re-reporting the case that the annotation file itself cites; curation from a non-PMID reference; and knowledge a tool encodes from sources other than phenotype.hpoa. Absence of detected overlap therefore lowers leakage risk without eliminating it. The flag is a measurable, recomputable indicator, not proof of independence.

Third, whether annotation overlap *actually* inflates a given tool’s measured accuracy is an empirical question this resource enables but does not answer: the benchmark supplies the cases, labels and flags needed to test it, and quantifying the effect for specific tools is left to future work.

Fourth, the protein-coding restriction excludes a small but clinically relevant class of causal genes. Both candidate-list variants draw distractors from the pinned HGNC protein-coding set, so a case whose causal gene lies outside it cannot be represented; three sampled neurological cases (two *RNU4-2*, one *RNU2-2*) were dropped for this reason (Section 2.1). Both are small nuclear RNA genes recently reported as causes of neurodevelopmental disorder [39,40], so this is a substantive scope restriction: the benchmark cannot measure how well a tool recovers non-protein-coding causal genes—an area where literature-based tools might hold an advantage over knowledge bases curated around protein-coding disease genes. Extending the candidate space is a natural direction for a future release.

Fifth, the hard candidate lists are not neutral across tool classes. We select distractors by Resnik similarity over genes_to_phenotype, the same annotation resource from which knowledge-base tools derive their gene–phenotype associations, so the hard variant is populated with the genes most confusable to a phenotype-similarity scorer; genes lacking HPO annotations can never be selected, and the hard *×* overlap-absent cell inherits both properties. We retain the variant because differential diagnosis against phenotype-similar competitors is the clinically realistic setting and the construction is recomputable. Read results on it as performance against an HPO-similarity-defined adversary: standard-versus-hard comparisons are informative within a tool, whereas comparisons between tool classes are more safely made on the standard variant.

Lastly, two disease classes are excluded by construction, so the cohort says nothing about them. Cases under the MONDO chromosomal-disorder (MONDO:0019042) and mitochondrial-disease (MONDO:0044970) roots were removed at the inclusion stage (Section 2.1). The mitochondrial exclusion is broader than the design strictly requires, because it also removes nuclear-encoded mitochondrial disease, which the 50-gene design could in principle represent. Users evaluating tools in these areas should not generalise from this cohort. Restoring nuclear-encoded mitochondrial disease would require replacing the disease-class exclusion with a per-case check on the causal gene’s genomic location, a straightforward extension for a future release.

## Ethics declarations

This work derives entirely from previously published, de-identified case data distributed in the GA4GH Phenopacket Store under CC BY 4.0, and from open ontology and literature resources. No new human-subjects research was conducted and no ethical approval was required.

## Author contributions

J.A. conceived and designed the resource (Conceptualization, Methodology), assembled the benchmark cohort and its metadata layers (Data curation), implemented the corpus, indexing, cohort-construction and validation pipelines (Software), performed the technical validation and the retrieval-substrate and clustering analyses (Formal analysis, Investigation, Validation), produced the figures (Visualization), secured the predoctoral funding that supported the work (Funding acquisition), and wrote the manuscript (Writing — original draft). H.E.M. and V.Y. supervised the work (Supervision), contributed to its conception and scope (Conceptualization), and revised the manuscript (Writing — review & editing). All authors read and approved the submitted version.

## Acknowledgements

This resource is built entirely on openly licensed community infrastructure. We thank the Monarch Initiative and the GA4GH Phenopacket Store curators, whose case-level curation makes a benchmark of this kind possible; the Human Phenotype Ontology, Mondo Disease Ontology and HGNC teams for maintaining the pinned vocabularies the cohort is built on; the National Library of Medicine for the PubMed Central Open Access subset; and the maintainers of Qdrant, fastembed and the PubMedBERT embedding model. We also acknowledge the clinicians, researchers and families whose published case reports underlie every case in this cohort.

During the preparation of this manuscript the authors used Claude Opus 5 (Anthropic) to assist with language editing and manuscript restructuring. All AI-assisted output was critically reviewed, verified and revised by the authors, who take full responsibility for the accuracy, originality and integrity of the final manuscript. Claude Opus 5 was additionally used to assist with code development and debugging; all AI-assisted code was reviewed, tested and validated by the authors. No AI system was used to determine cohort inclusion, ground-truth labels or scientific conclusions.

## Code availability

The construction and validation code (corpus parsing, index build, cohort pipeline stages, annotation-overlap and recency metadata generation, and checksum/manifest tooling) is archived under AGPL-3.0 at https://doi.org/10.6084/m9.figshare.32814491 (Data Citation 3). The deposit is the snapshot at release tag paper-methods-v1.4, which is the normative state of the code described here.

Standard-cohort regeneration corresponds to pipeline stages 13–20 (scripts/cases/); the hard-variant candidate lists are produced by scripts/cases/18b_build_hard_candidates.py; the annotation-overlap flag is produced by scripts/eval/compute_annotation_overlap.py; the case-clustering statistics by scripts/eval/compute_clustering_stats.py.

The retrieval-substrate characterisations reported in Section 4 are produced by scripts/eval/ validate_retrieval_substrate.py (source-article recall and symbol grounding) and the chunk-set fingerprint by scripts/corpus/compute_chunk_fingerprint.sh. All four figures, and the strict-exceedance decomposition quoted in Section 4, regenerate from the released cohort and the pinned HPO release via scripts/manuscript/render_p1_figures.py.

The retrieval substrate is built by scripts/corpus/02_extract_and_parse_ftp.py (tarball extraction, JATS parsing, retraction exclusion), scripts/corpus/03_normalize_dedupe_filter. py (deduplication, genetics-relevance filter, schema normalisation), scripts/corpus/04_chunk_ normalized.py (section-aware chunking and chunk identifiers), scripts/embedding/05_embed_ chunks.py (dense and sparse vectors), and scripts/indexing/10_create_qdrant_index.py plus scripts/indexing/06 _upload_to_qdrant.py (collection schema and upload). Scripts 01_demo_fetch_pmc.py, 06_parse_jats_xml.py, 07_filter_corpus.py and 08_section_ aware_chunking.py implement a small demonstration path over an E-utilities-fetched sample and are not part of the released build.

